# JuPOETs: A Constrained Multiobjective Optimization Approach to Estimate Biochemical Model Ensembles in the Julia Programming Language

**DOI:** 10.1101/056044

**Authors:** David Bassen, Michael Vilkhovoy, Mason Minot, Jonathan T Butcher, Jeffrey D. Varner

**Author notes:** Corresponding author: Jeffrey D. Varner, Professor, School of Chemical and Biomolecular Engineering, 244 Olin Hall, Cornell University, Ithaca NY, 14853.

## Abstract

Ensemble modeling is a well established approach for obtaining robust predictions and for simulating course grained population behavior in deterministic mathematical models. In this study, we present a multiobjective based technique to estimate model ensembles, the Pareto Optimal Ensemble Technique in the Julia programming language (JuPOETs). JuPOETs integrates simulated annealing with Pareto optimality to estimate parameter ensembles on or near the optimal tradeoff surface between competing training objectives. We demonstrated JuPOETs on a suite of multiobjective problems, including test functions with parameter bounds and system constraints as well as for the identification of a proof-of-concept biochemical model with four conflicting training objectives. JuPOETs identified optimal or near optimal solutions approximately six-fold faster than a corresponding implementation in Octave for the suite of test functions. For the proof-of-concept biochemical model, JuPOETs produced an ensemble of parameters that gave both the mean of the training data for conflicting data sets, while simultaneously estimating parameter sets that performed well on each of the individual objective functions. JuPOETs can be adapted to solve many problem types, including mixed binary and continuous variable types, bilevel optimization problems and constrained problems without altering the base algorithm. JuPOETs can be installed using the Julia package manager from the JuPOETs GitHub repository at https://github.com/varnerlab/POETs.jl.

## Background

Ensemble modeling is a well established approach for obtaining robust predictions and course grained population behavior in large deterministic mathematical models. It is often not possible to uniquely identify all the parameters in biochemical models, even when given extensive training data [1]. Thus, despite significant advances in standardizing biochemical model identification [2], the problem of estimating model parameters from experimental data remains challenging. Ensemble approaches address parameter uncertainty in systems biology and other fields like weather prediction [3–6] by using parameter families instead of single best-fit parameter sets. Parameter families can be selected based upon simulation error, along with other criterion such as diversity or steady-state peformance. Simulations using parameter ensembles can estimate confidence intervals on model variables, and robustly constrain model predictions, despite having many poorly constrained parameters [7, 8]. There are many techniques to generate parameter ensembles. Battogtokh et al., Brown et al., and later Tasseff et al. generated experimentally constrained parameter ensembles using a Metropolis-type random walk [3, 5, 9, 10]. Liao and coworkers developed methods that generate ensembles that all approach the same steady-state, for example one determined by fluxomics measurements [11]. They have used this approach for model reduction [12], strain engineering [13, 14] and to study the robustness of non-native pathways and network failure [15]. Maranas and coworkers have also applied this method to develop a comprehensive kinetic model of bacterial central carbon metabolism, including mutant data [16]. We and others have used ensemble approaches, generated using both sampling and optimization techniques, to robustly simulate a wide variety of signal transduction processes [9, 10, 17–19], neutrophil trafficking in sepsis [20], patient specific coagulation behavior [21] and to capture cell to cell variation [22]. Thus, ensemble approaches are widely used to robustly simulate a variety of biochemical systems.

Identification of biochemical models with hundreds or even thousands of states and parameters may not be tractable as a single objective optimization problem. Further, large models require significant training data perhaps taken from diverse sources, for example different laboratories or cell-lines. These data are often heterogenous, and contain intrinsic conflicts that complicate parameter estimation. Parameter ensembles which optimally balance tradeoffs between submodels and conflicts in training data can lead to robust model performance. Multiobjective optimization is an ensemble generation technique that naturally balances conflicting training data. Previously, we developed the Pareto Optimal Ensemble Technique (POETs) algorithm to address the challenge of competing or conflicting objectives. POETs, which integrates simulated annealing (SA) and multiobjective optimization through the notion of Pareto rank, estimates parameter ensembles which optimally trade-off between competing (and potentially conflicting) experimental objectives [23]. However, the previous implementation of POETs, in the Octave programming language [24], suffered from poor performance and was not configurable. For example, Octave-POETs does not accommodate user definable objective functions, bounds and problem constraints, cooling schedules, different variable types e.g., a mixture of binary and continuous design variables or custom diversity generation routines. Octave-POETs was also not well integrated into a package or source code management (SCM) system. Thus, upgrades to the approach containing new features, or bug fixes were not centrally managed.

## Implementation

In this study, we present an open-source implementation of the Pareto optimal ensemble technique in the Julia programming language (JuPOETs). JuPOETs offers many advantages and improvements compared to Octave-POETs. JuPOETs takes advantage of the unique features of Julia. Julia is a cross-platform, high-performance programming language for technical computing that has performance comparable to C but with syntax similar to MATLAB/Octave and Python [25]. Julia also offers a sophisticated compiler, distributed parallel execution, numerical accuracy, and an extensive function library. Further, the architecture of JuPOETs takes advantage of the first-class function type in Julia allowing user definable behavior for all key aspects of the algorithm, including objective functions, custom diversity generation logic, linear/non-linear parameter constraints (and parameter bounds constraints) as well as custom cooling schedules. Julia’s ability to naturally call other languages such as Python or C also allows JuPOETS to be used with models implemented in a variety of languages across many platforms. Additionally, Julia offers a built-in package manager which is directly integrated with GitHub, a popular web-based Git repository hosting service offering distributed revision control and source code management. Thus, JuPOETs can be adapted to many problem types, including mixed binary and continuous variable types, bilevel problems and constrained problems without altering the base algorithm, as was required in the previous POETs implementation.

**JuPOETs optimization problem formulation**. JuPOETs solves the *κ*-dimensional constrained multiobjective optimization problem:

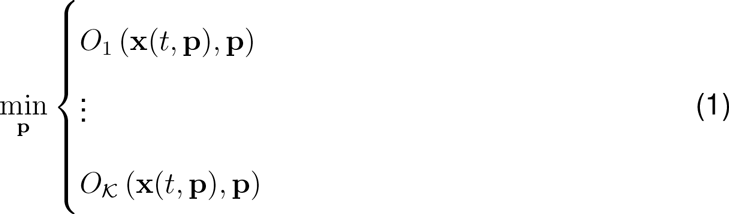

subject to:

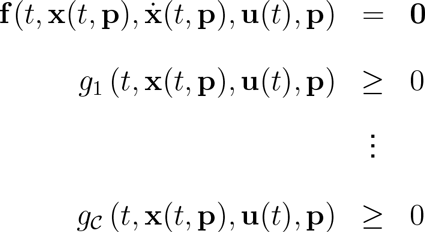

and parameter bound constraints:

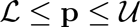

using a modified simulated annealing approach. The quantity *t* denotes time, x(*t*, p) denotes the model state (with an initial state x_0_), and u(*t*) denotes an input vector. The terms 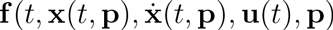 denote the system of model equations (e.g., differential equations, differential algebraic equations or linear/non-linear algebraic equations) where p denotes the unknown parameter vector (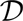 × 1). The parameter search can be subject to parameter bound constraints, where 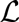 and 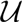 denote the lower and upper parameter bounds, respectively as well as *C* problem specific constraints *g_i_*(*t*,x(*t*,p),u(*t*),p), *i* = 1,…, *C*.

JuPOETs integrates simulated annealing with Pareto optimality to estimate parameter sets on or near the optimal tradeoff surface between competing training objectives (Fig. 1 and Algorithm 1). The central idea of POETs is a mapping between the value of the objective vector evaluated at p_*i*+1_ (parameter guess at iteration *i* + 1) and Pareto rank. JuPOETs calculates the performance of a candidate parameter set p_*i*+1_ by calling the user defined `objective` function; `objective` takes a parameter set as an input and returns a *κ* × 1 objective vector. Candidate parameter sets are generated by the user supplied `neighbor` function. The error vector associated with p_*i*+1_ is ranked using the builtin Pareto `rank` function, by comparing the current error at iteration *i* + 1 to the error archive 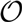_*i*_ (all error vectors up to iteration *i* − 1 meeting a ranking criteria). Pareto rank is a measure of distance from the trade-off surface; parameter sets on or near the optimal trade-off surface between the objectives have a rank equal to 0 (no other current parameter sets are better). Sets with increasing non-zero rank are progressively further away from the optimal trade-off surface. Thus, a parameter set with a rank = 0 is *better* in a trade-off sense than rank > 0. We implemented the Fonseca and Fleming ranking scheme in the builtin `rank` function [26]:

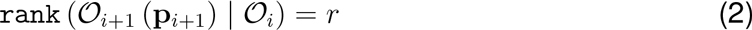

where rank *r* is the number of parameter sets that dominate (are better than) parameter set p_*i*+1_, and 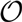_*i*+1_ (p_*i*+1_) denotes the objective vector evaluated at p_*i*+1_. We used the Pareto rank to inform the SA calculation. The parameter set p_*i*+1_ was accepted or rejected by the SA, by calculating an acceptance probability 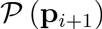:

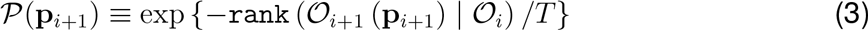

where *T* is the computational annealing temperature. As `rank` 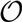_*i*+1_ (p_*i*+1_) | 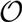_*i*_) ⟶ 0, the acceptance probability moves toward one, ensuring that we explore parameter sets along the Pareto surface. Occasionally, (depending upon *T*) a parameter set with a high Pareto rank was accepted by the SA allowing a more diverse search of the parameter space. However, as *T* is reduced, the probability of accepting a high-rank set occurring decreases. Parameter sets could also be accepted by the SA but not permanently archived in *S_i_*. Only parameter sets with rank less than or equal to threshold (rank ≤4 by default) were included in *S_i_*, where the archive was re-ranked and filtered after every new parameter set was accepted. Parameter bounds were implemented in the `neighbor` function as box constraints, while problem specific constraints were implemented in `objective` using a penalty method:

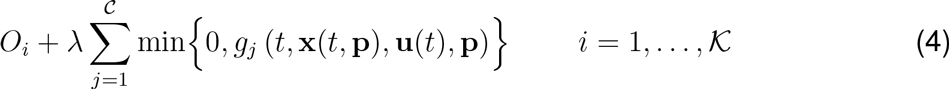

where λ denotes the penalty parameter (λ = 100 by default). However, because both the neighbor and objective functions are user defined, different constraint implementations are easily defined. JuPOETs can be installed using the Julia package manager from the JuPOETs repository at https://github.com/varnerlab/POETs.jl. Sample code is included in the `sample/biochemical` subdirectory of the JuPOETs repository to help users get started using JuPOETs in their projects.

**Fig. 1:**
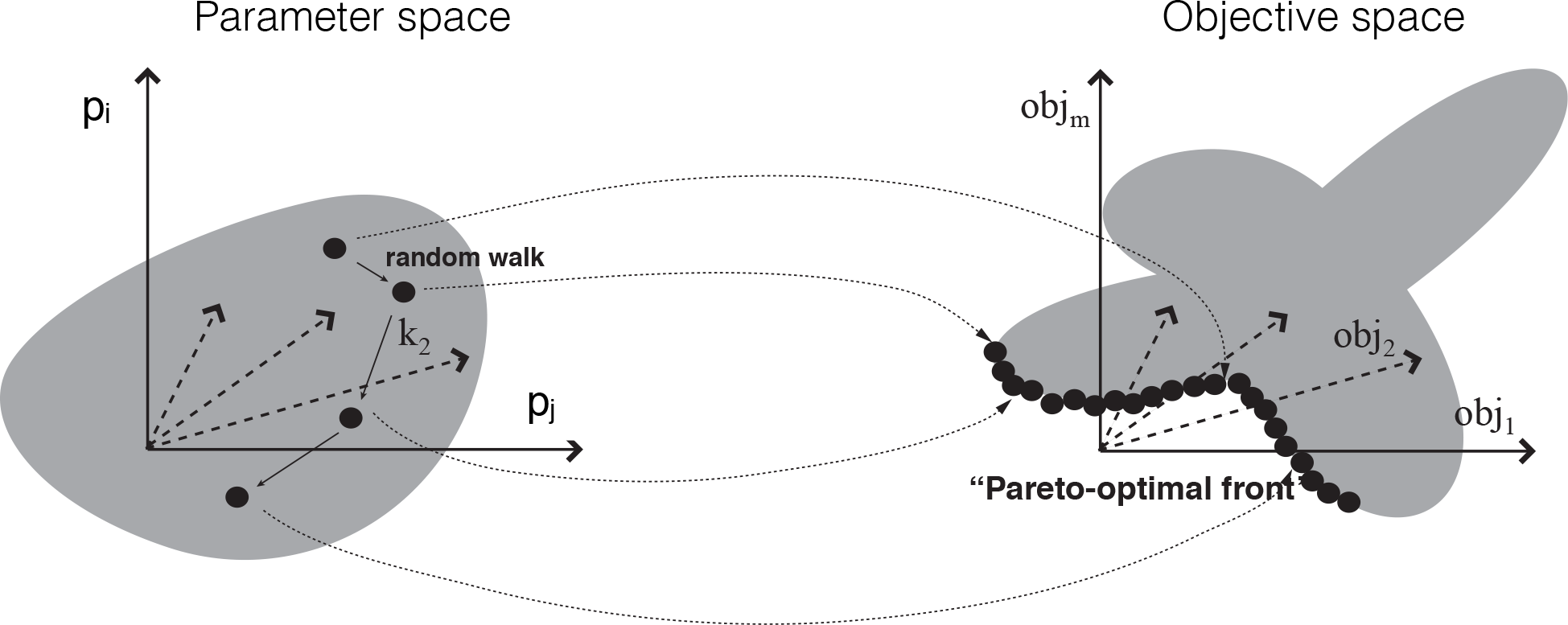
Schematic of multiobjective parameter mapping. The performance of any given parameter set is mapped into an objective space using a ranking function which quantifies the quality of the parameters. The distance away from the optimal tradeoff surface is quantified using the Pareto ranking scheme of Fonseca and Fleming in JuPOETs.

## Results and Discussion

JuPOETs identified optimal or nearly optimal solutions significantly faster than Octave-POETs for a suite of multiobjective test problems (Table 1). The wall-clock time for JuPOETs and Octave-POETs was measured for 10 independent trials for each of the test problems. The same `cooling, neighbor, acceptance,` and `objective` logic was employed between the implementations, and all other parameters were held constant. For each test function, the search domain was partitioned into 10 segments, where an initial parameter guess was drawn from each partition. The number of search steps for each temperate was 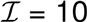 for all cases, and the cooling parameter was *α* = 0.9. On average, JuPOETs identified optimal or near optimal solutions for the suite of test problems six-fold faster (60s versus 400s) than Octave-POETs (Fig. 2). JuPOETs produced the characteristic tradeoff curves for each test problem, given both parameter bound and problem constraints (Fig. 3). Thus, JuPOETs estimated an ensemble of solutions to constrained multiobjective optimization test problems significantly faster than the current Octave implementation. Next, we tested JuPOETs on a proof-of-concept biochemical model identification problem.

**Fig. 2:**
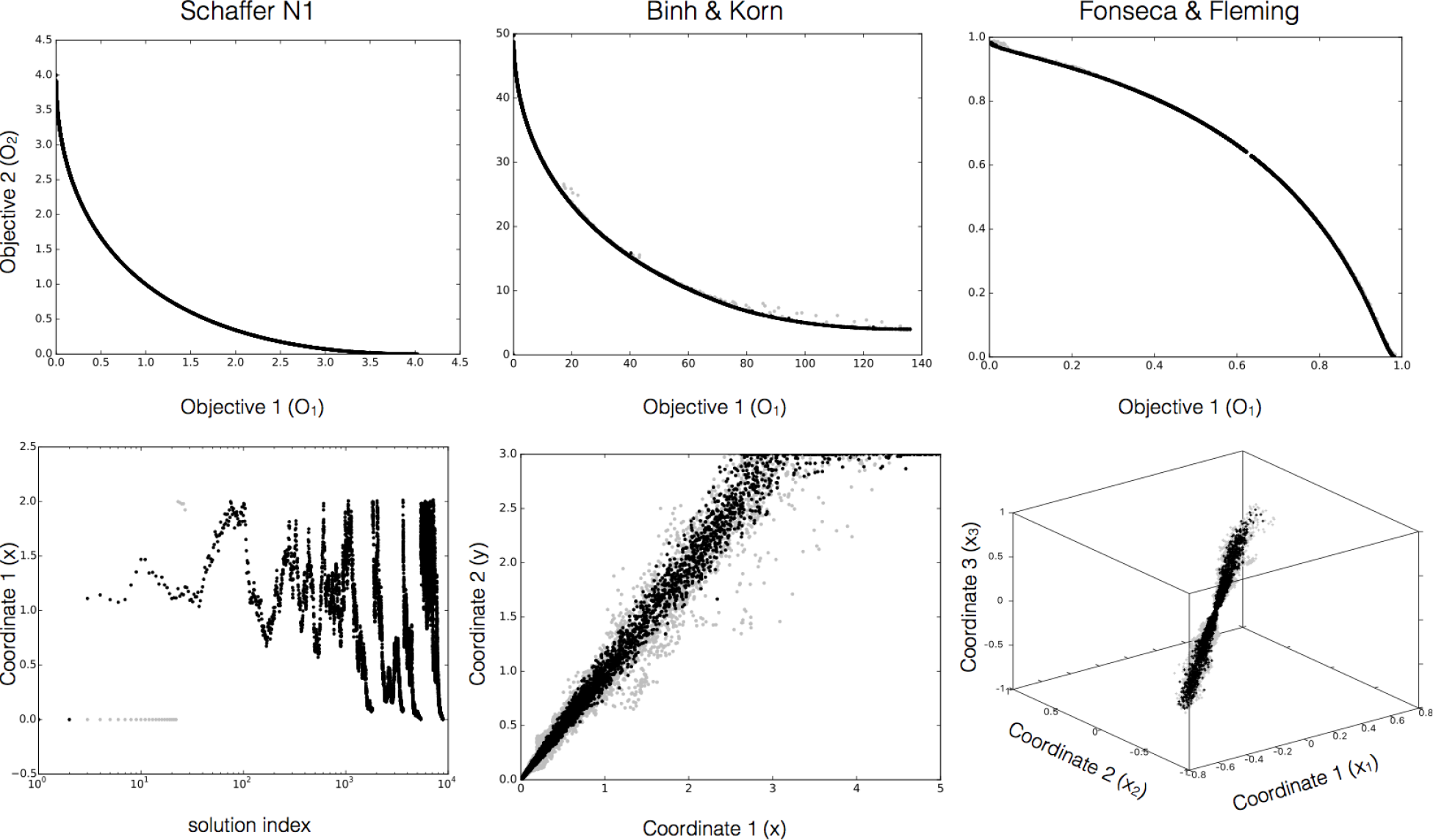
The performance of JuPOETs on the multi-objective test suite. The execution time (wall-clock) for JuPOETs and POETs implemented in Octave was measured for 10 independent trials for the suite of test problems. The number of steps per temperature 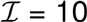, and the cooling parameter *α* = 0.9 for all cases. The problem domain was partitioned into 10 equal segments, an initial guess was drawn from each segment. For each of the test functions, JuPOETs estimated solutions on (rank zero solutions, black) or near (gray) the optimal tradeoff surface, subject to bounds and problem constraints.

**Fig. 3:**
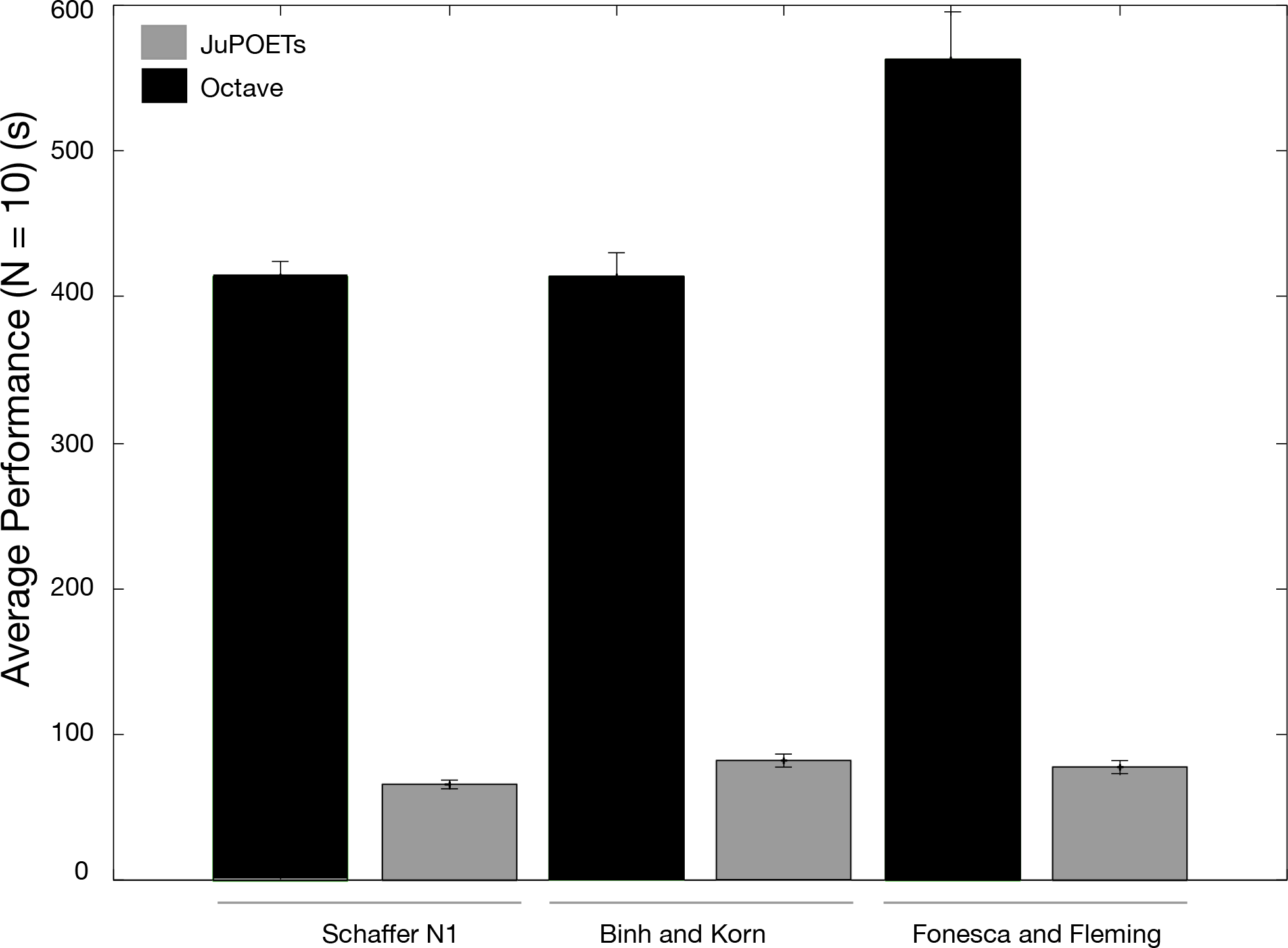
Representative JuPOETs solutions for problems in the multi-objective test suite. The number of steps per temperature 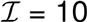, and the cooling parameter *α* = 0.9 for all cases. The problem domain was partitioned into 10 equal segments, an initial guess was drawn from each segment. For each of the test functions, JuPOETs estimated solutions on (rank zero solutions, black) or near (gray) the optimal tradeoff surface, subject to bounds and problem constraints.

**Table 1:** Multi-objective optimization test problems. We tested the JuPOETs implementation on three twodimensional test problems, with one-, two- and three-dimensional parameter vectors. Each problem had parameter bounds constraints, however, on the Binh and Korn function had additional non-linear problem constraints. For the Fonesca and Fleming problem, N = 3.

| Name | Dimension | Function | Domain | Constraints |
| --- | --- | --- | --- | --- |
| Schaffer function | 1 | $O_1(x) = x^2$<br>$O_2(x) = (x - 2)^2$ | $-10 \leq x \leq 10$ | |
| Binh and Korn function | 2 | $O_1(x, y) = 4x^2 + 4y^2$<br>$O_2(x, y) = (x - 5)^2 + (y - 5)^2$ | $0 \leq x \leq 5$<br>$0 \leq y \leq 3$ | $g_1(x, y) = (x - 5)^2 + y^2 \leq 25$<br>$g_2(x, y) = (x - 8)^2 + (y + 3)^2 \geq 7.7$ |
| Fonseca and Fleming function | 3 | $O_1(x_i) = 1 - \exp\left(-\sum_{i=1}^N \left(x_i - \frac{1}{\sqrt{N}}\right)^2\right)$<br>$O_2(x_i) = 1 - \exp\left(-\sum_{i=1}^N \left(x_i + \frac{1}{\sqrt{N}}\right)^2\right)$ | $-4 \leq x_i \leq 4$ | |

JuPOETs estimated an ensemble of biochemical models that was consistent with the mean of synthetic training data (Fig. 4). Four synthetic training data sets were generated from a prototypical biochemical network consisting of 6 metabolites and 7 reactions (Fig. 4, inset right). We considered a common case in which the same measurements were made on four hypothetical cell types, each having the same biological connectivity but different performance. Network dynamics were modeled using the hybrid cybernetic model with elementary modes (HCM-EM) approach of Ramkrishna and coworkers [27]. In the HCM-EM approach, metabolic networks are first decomposed into a set of elementary modes (EMs) (chemically balanced steady-state pathways, see [28]). Dynamic combinations of elementary modes are then used to characterize network behavior. Each elementary mode is catalyzed by a pseudo enzyme; thus, each mode has both kinetic and enzyme synthesis parameters. The proof of concept network generated 6 EMs, resulting in 13 model parameters. The synthetic data was generated by randomly varying these parameters. JuPOETs produced an ensemble of approximately dim *S* ≃ 13,000 parameters that captured the mean of the measured data sets for extracellular metabolites and cellmass (Fig. 4A and B). JuPOETs minimized the difference between the simulated and measured values for A_*e*_, B_*e*_, C_*e*_ and cellmass, where the residual for each data set was treated as a single objective (leading to four objectives). The 95% confidence estimate produced by the ensemble was consistent with the mean of the measured data, despite having significant uncertainty in the training data. JuPOETs produced a consensus estimate of the synthetic data by calculating optimal trade-offs between the training data sets (Fig. 4C). Multiple trade-off fronts were visible in the objective plots, for example between data set 3 (O_3_) and data set 2 (O_2_). Thus, without a multiobjective approach, it would be challenging to capture these data sets as fitting one leads to decreased performance on the other. However, the ensemble contained parameter sets that described each data set independently (Fig. 5). Thus, JuPOETs produced an ensemble of parameters that gave the mean of the training data for conflicting data sets, while simultaneously estimating parameter sets that performed well on each individual objective function.

**Fig. 4:**
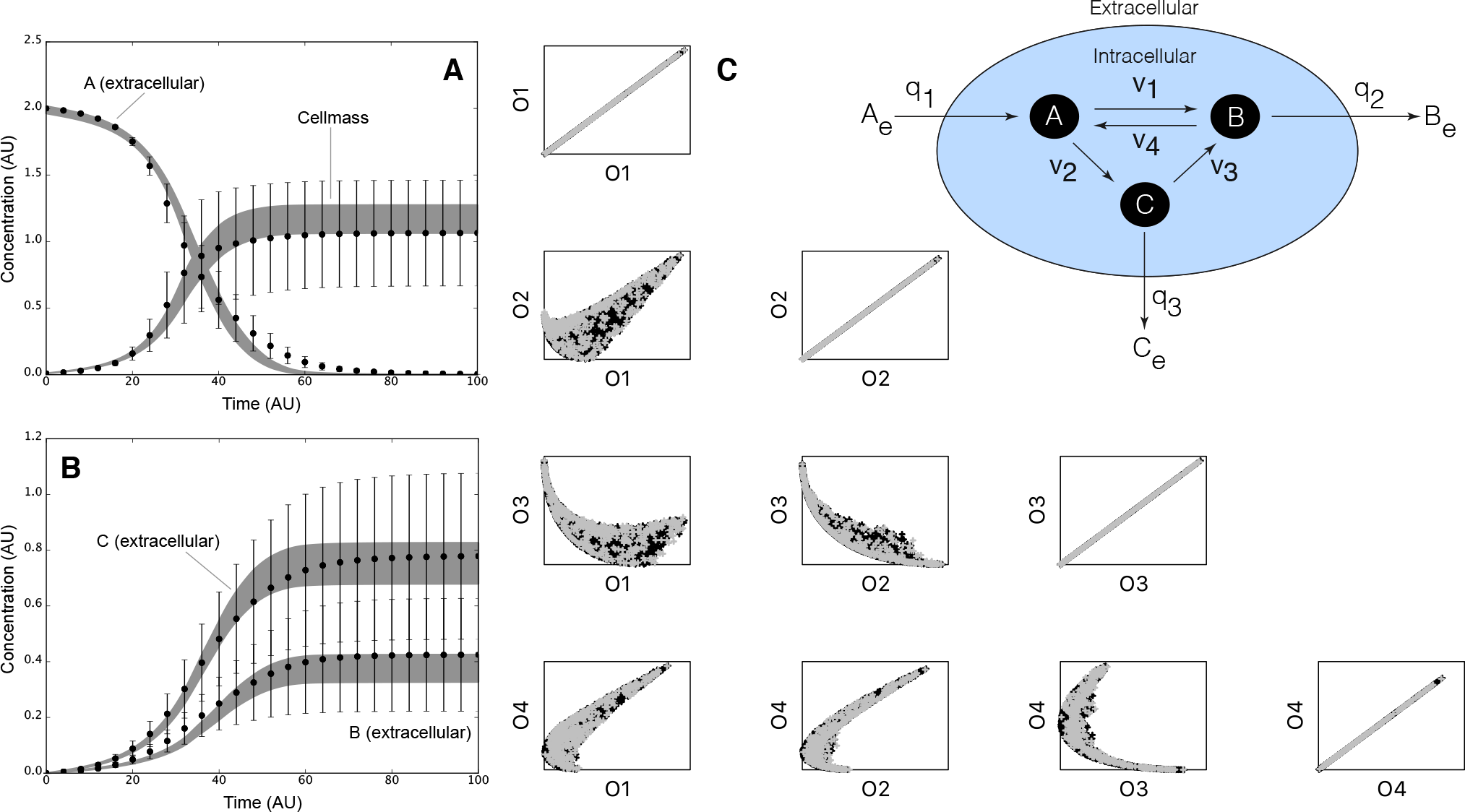
Proof of concept biochemical network study. Inset right: Prototypical biochemical network with six metabolites and seven reactions modeled using the hybrid cybernetic approach (HCM). Intracellular cellmass precursors *A*, *B*, and *C* are balanced (no accumulation) while the extracellular metabolites *A_e_*, *B_e_*, and *C_e_* are dynamic. The oval denotes the cell boundary, *q_j_* is the *j*th flux across the boundary, and *v_k_* denotes the *k*th intracellular flux. Four data sets (each with *A_e_*,*B_e_*,*C_e_* and cellmass measurements) were generated by varying the kinetic constants for each biochemical mode. Each data set was a single objective in the JuPOETs procedure. A: Ensemble simulation of extracellular substrate *A_e_* and cellmass versus time. B: Ensemble simulation of extracellular substrate *B_e_* and *C_e_* versus time. The gray region denotes the 95% confidence estimate of the mean ensemble simulation. The data points denote mean synthetic measurements, while the error bars denote the 95% confidence estimate of the measurement computed over the four training data sets. C: Trade-off plots between the four training objectives. The quantity *O_j_* denotes the jth training objective. Each point represents a member of the parameter ensemble, where black denotes a rank 0 set, while gray denotes rank 1 set.

**Fig. 5:**
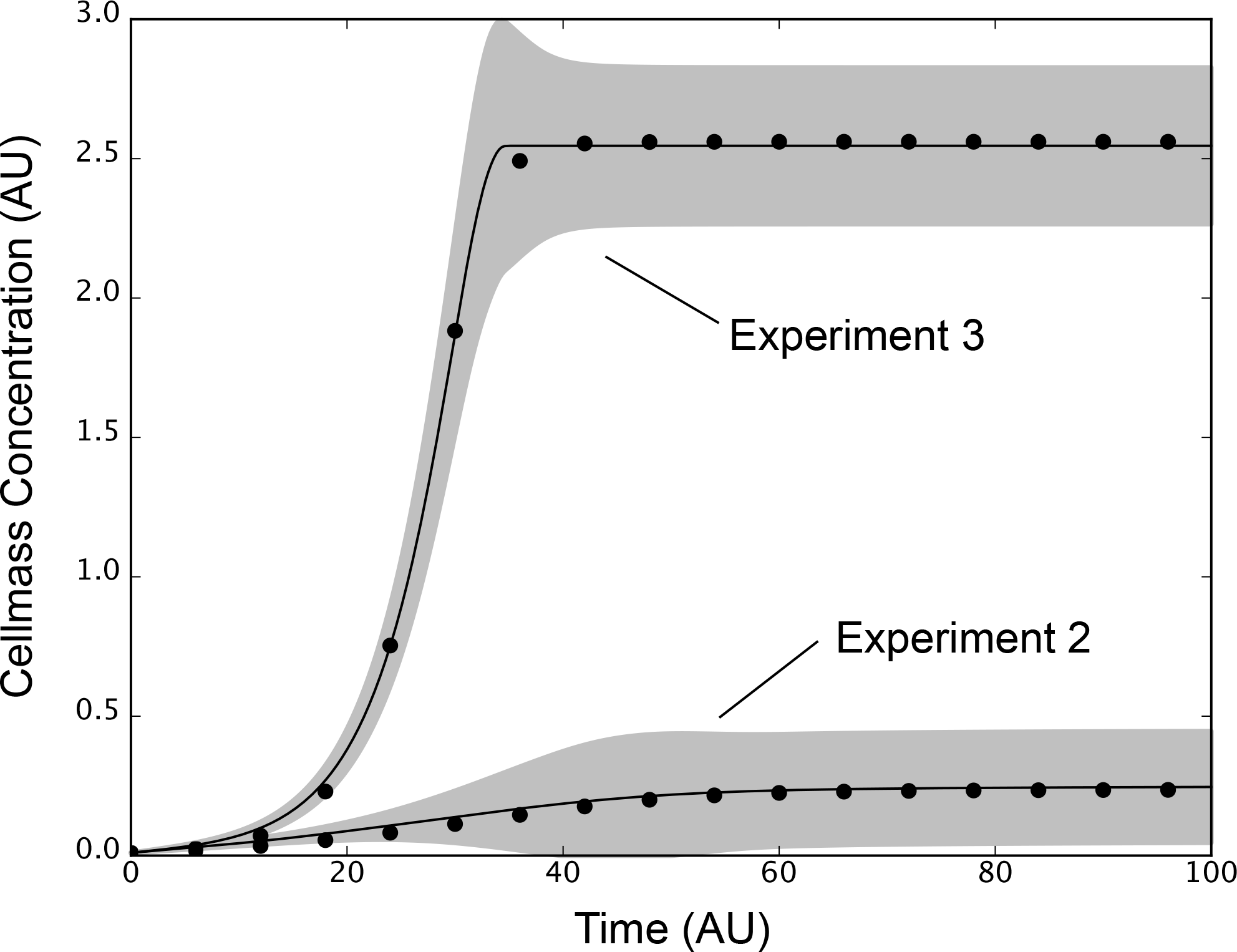
Experiment to experiment variation is captured by a single ensemble. Cellmass measurements (points) versus time for experiment 2 and 3 were compared with ensemble simulations. The full ensemble was sorted by simultaneously selecting the top 25% of solutions for each objective with rank ≤ 1. The best fit solution for each objective (line) ± 1-standard deviation (gray region) for experiment 2 and 3 brackets the training data despite significant differences the training values between the two data sets.

## Conclusions

JuPOETs is a significant advance over the previous POETs implementation. It offers improved performance and is highly adaptable to different problem types. We demonstrated JuPOETs on a suite of test problems, and a proof-of-concept biochemical model. However, there are several areas that could be explored further to improve JuPOETs. First, JuPOETs should be compared with other multiobjective evolutionary algorithms (MOEAs) to determine its relative performance on test and real world problems. Many evolutionary approaches e.g., the nondominated sorting genetic algorithm (NSGA) family of algorithms, have been adapted to solve multiobjective optimization problems [29, 30]. It is unclear if JuPOETs will perform as well as these other approaches; one potential advantage that JuPOETs may have is the local refinement step which temporarily reduces the problem to a single objective formulation. Previously, this hybrid approach led to better convergence on a proof-of-concept signal transduction model [23]. For many real world parameter estimation problems, the bulk of the execution time is spent evaluating the objective functions. One strategy to improve performance could be to optimize surrogates [31], while another would be parallel execution of the objective functions. Currently, JuPOETs serially evaluates the objective function vector. However, parallel evaluation of the objective functions could be easily implemented using a variety of techniques without changing the main run loop of JuPOETs. Because of the flexible function pointer architecture of JuPOETs, the only changes required are in the user defined objective function. Taken together, JuPOETs has demonstrated improved flexibility, and performance over POETs in parameter identification and ensemble generation for multiple objectives. JuPOETs has the potential for widespread use due to the flexibility of the implementation, and the high level syntax and distribution tools native to Julia.

## Competing interests

The authors declare that they have no competing interests.

## Funding

This study was supported by an award from the National Science Foundation (NSF CBET-0955172) and the National Institutes of Health (NIH HL110328) to J.B, and by a National Science Foundation Graduate Research Fellowship (DGE-1144153) to D.B.

## Author’s contributions

J.V developed the software presented in this study. M.M and M.V developed the proof-of-concept biochemical model. The manuscript was prepared and edited for publication byD.B, J.B. and J.V.

---

**input**: User specified neighbor, objective, acceptance and cooling functions. Initial parameter guess (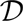 × 1)

**Output**: Rank archive 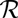, parameter solution archive *S* and objective archive 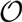

**Algorithm 1:**
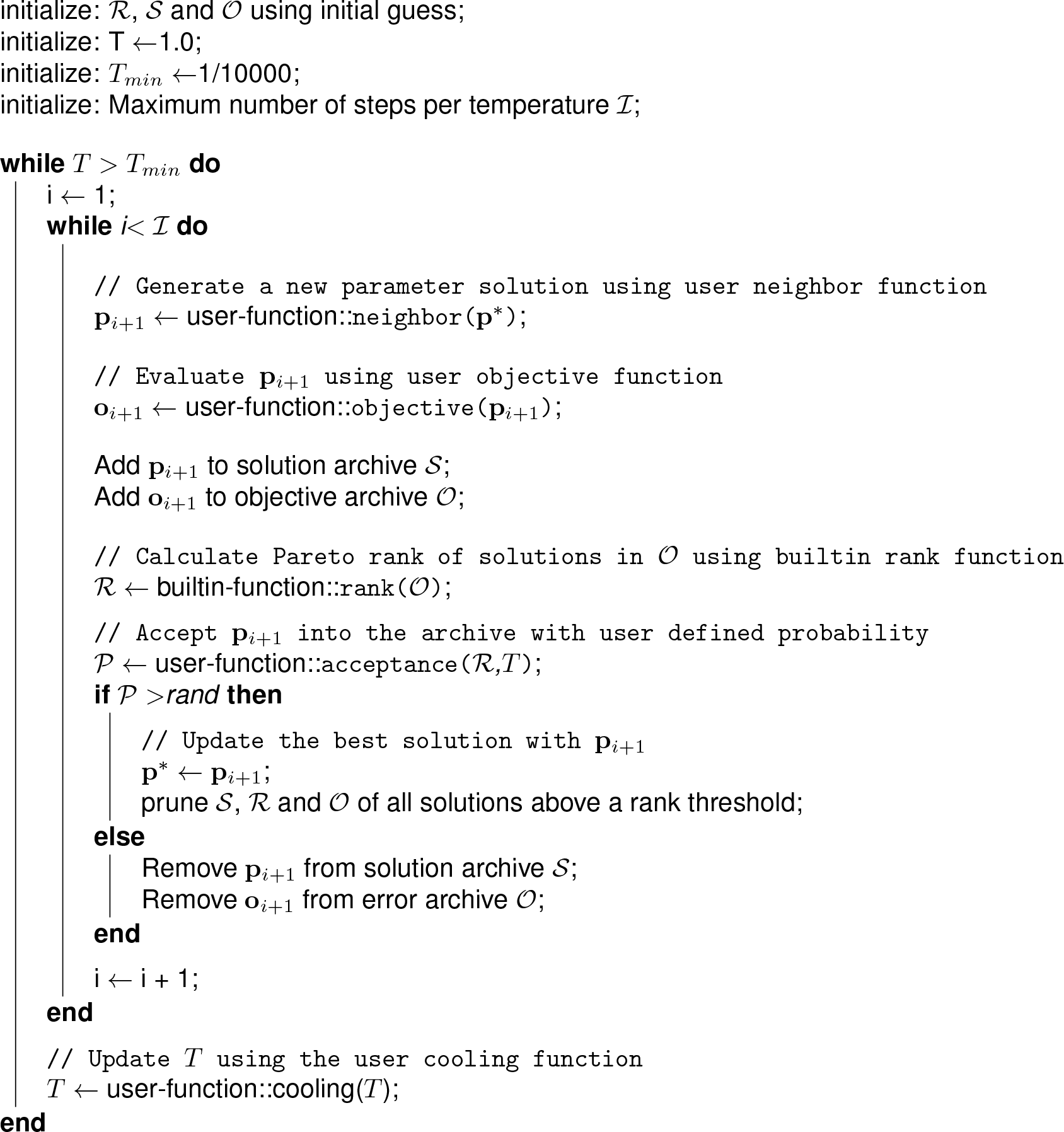
Pseudo-code for the main run-loop of JuPOETs. The user specifies the `neighbor, acceptance, cooling` and `objective` functions along with an initial parameter guess. The rank archive 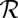, solution archive *S* and objective archive 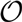 are initialized from the initial guess. The initial guess is perturbed in the `neighbor` function, which generates a new solution whose performance is evaluated using the user supplied `objective` function. The new solution and objective values are then added to the respective archives and ranked using the builtin `rank` function. If the new solution is accepted (based upon a probability calculated with the user supplied `acceptance` function) it is added to the solution and objective archive. This solution is then perturbed during the next iteration of the algorithm. However, if the solution is not accepted, it is removed from the archive and discarded. The computational temperature is adjusted using the user supplied `cooling` function after each 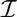 iterations.

